# SACSANN: identifying sequence-based determinants of chromosomal compartments

**DOI:** 10.1101/2020.10.06.328039

**Authors:** Julie A Prost, Christopher JF Cameron, Mathieu Blanchette

## Abstract

Genomic organization is critical for proper gene regulation and based on a hierarchical model, where chromosomes are segmented into megabase-sized, cell-type-specific transcriptionally active (A) and inactive (B) compartments. Here, we describe SACSANN, a machine learning pipeline consisting of stacked artificial neural networks that predicts compartment annotation solely from genomic sequence-based features such as predicted transcription factor binding sites and transposable elements. SACSANN provides accurate and cell-type specific compartment predictions, while identifying key genomic sequence determinants that associate with A/B compartments. Models are shown to be largely transferable across analogous human and mouse cell types. By enabling the study of chromosome compartmentalization in species for which no Hi-C data is available, SACSANN paves the way toward the study of 3D genome evolution. SACSANN is publicly available on GitHub: https://github.com/BlanchetteLab/SACSANN

## Introduction

High-throughput chromosome conformation capture (Hi-C) provides a population estimate of intra- and inter-chromosomal Interaction Frequencies (IF) for all loci pairs of a genome (1). The output of a Hi-C experiment is typically stored in a set of intra- and inter-chromosomal IF matrices, whose rows and columns correspond to genomic bins of a fixed size (e.g., 50 kb). Lieberman-Aiden et al. (2009) (1) discovered mammalian genomes are segmented into two types of megabase-sized compartments: i) A(ctive) compartments, which have been linked to euchromatin and are gene rich, transcriptionally active regions; and ii) B (inactive) compartments, associated with heterochromatin and repressed gene expression. Contacts between genomic regions belonging to the same compartment are generally more frequent than those involving pairs of regions from different compartments, resulting in the typical “plaid pattern” seen across Hi-C IF matrices. Compartments have also been shown to be composed of one or more Topologically Associating Domains (TAD) (2), which are genomic regions whose loci preferentially interact with each other. Compartment annotations can be obtained by a simple Principal Component Analysis (PCA) (3) of Hi-C IF matrices, where the sign of the projection onto the first principal component divides the genome into A and B compartments (1).

A compartments are known to have high GC content and be enriched in activating chromatin marks (e.g., H3K27ac and H3K36me3) (1). Dixon et al. (2015) (4) further highlighted that compartments are cell-type specific and variable across cellular differentiation, with about 10% of loci switching compartment during the differentiation of human Embryonic Stem Cells (hESC) into Neuron Progenitors Cells (NPC). Overall, up to 36% of all compartments were altered during the differentiation of hESCs into four distinct cell types (namely NPCs, mesendoderm, mesenchymal and trophoblast-like cells). Moreover, these compartment alterations were found to correlate with corresponding changes in gene expression (4). Therefore, compartments are believed to play a role in cell-type specific gene expression profiles. In addition, Rao et al. (2014) (5) demonstrated that A/B compartments in human cell lines can be further partitioned into six types of sub-compartments (A1, A2, B1, B2, B3 and B4), each having its own genomic and epigenomic characteristics. These A/B sub-compartments and their characteristics have been suggested to be partially explained by cell-type specific transcription factor spatial networks (6).

The specific determinants of A/B compartment formation remain unclear. Fortin et al. (2015) (7) were able to apply eigenvector analysis to matrices of epigenetic data correlations and reconstruct compartments. Previous machine learning studies (8) have demonstrated that A/B compartments can be accurately predicted from ChIP-seq data using artificial neural networks. However, both of these methods are dependent on the availability of biochemical data for a given species and cell type, and provide little insight into the sequence encoding of genomic compartmentalization.

In this paper, we describe the supervised machine learning problem of classifying genomic loci into cell-type-specific A/B compartments using genomic sequence alone. We hypothesize that machine learning models limited to genomic sequence (or features derived from) as input will reveal sequence determinants of A/B compartment formation. While epigenetic modifications play an important role in establishing compartments (7, 8), these marks are deposited in a sequence-dependent manner. Therefore, the compartment structure in a given cell type is ultimately sequence-encoded. Although the problem of sequence-based compartment prediction has never been addressed, other aspects of 3D chromatin architecture prediction have been considered. Nikumbh and Pfeifer (2017) (9) demonstrated that long-range chromatin interactions can be predicted using a genomic sequence-based support vector machine. Whalen et al. (2016) (10) also showed that candidate enhancers and promoters genetic sequences could be used to predict enhancer-promoters interactions with ensemble boosted trees. These results are encouraging for the use of sequence features as the only inputs to an A/B compartment annotator.

Here, we describe a stacked artificial neural network model approach to predicting A/B compartments from genomic sequenc features called ‘Sequence-based Annotator of Chromosomal Compartments by Stacked Artificial Neural Networks’ (SACSANN). SACSANN takes as input features derived solely from the genomic DNA sequence of a given species, enabling compartment annotations for genomes where only the DNA sequence is available. Our cell-type specific models achieve high prediction accuracy on both hESC and mouse Embryonic Stem Cells (mESC), as well as the cell types of a mouse neuronal differentiation (mESC, NPC, and Cortical Neurons [CN]). Trained models are shown to be transferable between the investigated species, where SACSANN learns a set of rules to annotate A/B compartments from sequence that is applicable to both human and mouse genomes. In addition, we investigate SACSANN’s input features to gain further insights into the underlying sequence determinants and evolutionary processes that may impact A/B compartment formation.

## Results

### A/B compartments can be predicted from sequence-level features

We developed SACSANN to predict A/B compartments using features engineered from a reference genome sequence (Fig. 1). The sequence-level features needed to train the model are derived from partitioning the genome into 100 kb bins and counting the occurrences of specific computationally-identified genetic elements in each bin. These include the GC-content as well as computationally predicted binding sites for 334 transcription factors in human and mouse and 41 (35) families of Transposable Elements (TE) in human (mouse). We first perform a cell-type specific feature-selection by retaining the 100 features with the highest feature importance, based on a random forest predictor. We then train two stacked artificial neural networks that classify each input vector as belonging to either an A or B compartment. SACSANN is trained and evaluated using chromosome-wise leave-one-out cross-validation.

**Fig. 1.**
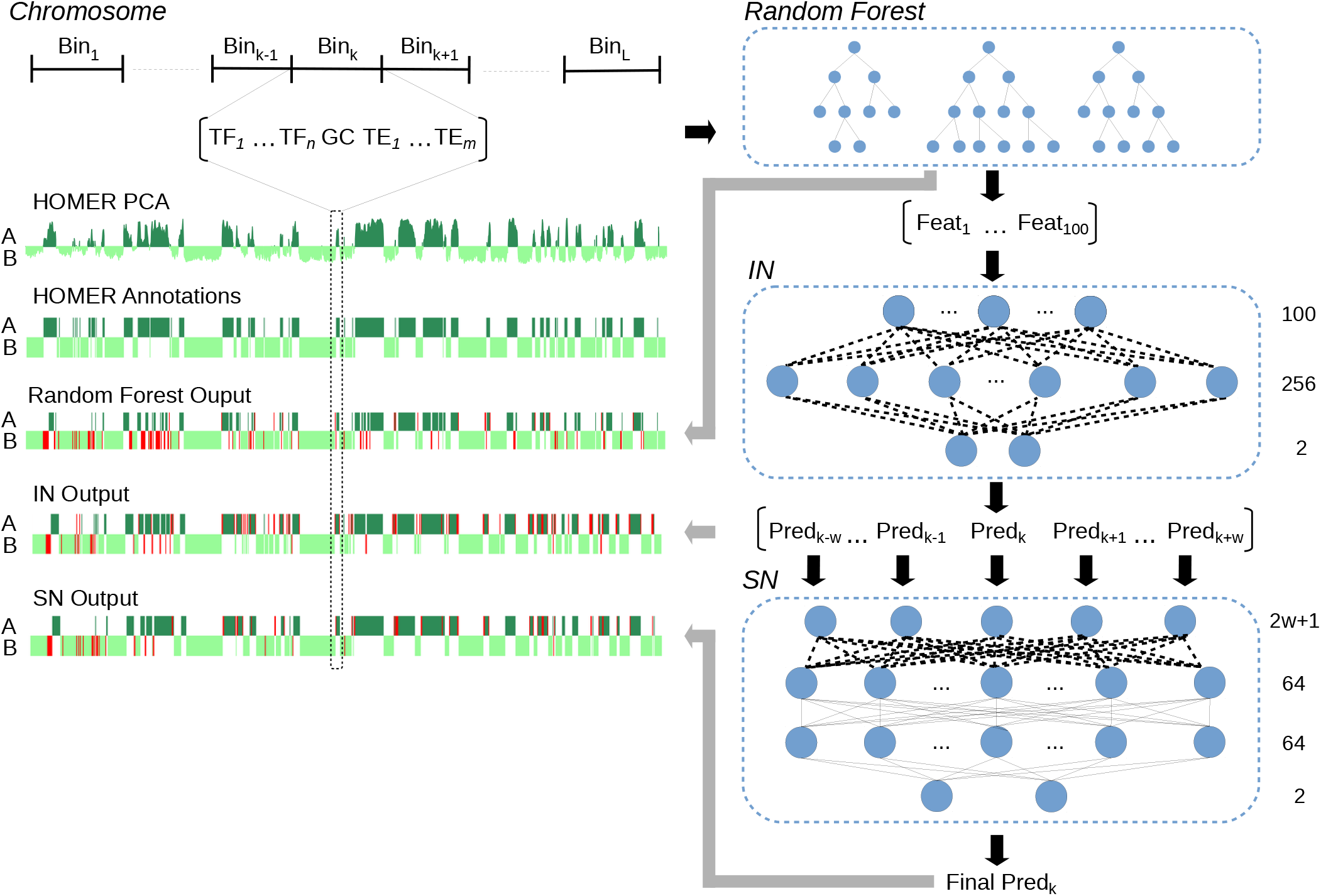
Summary of SACSANN compartment prediction pipeline. Each chromosome is first divided into fixed-sized bins (i.e., 100 kb). The sequence of each bin is analyzed to obtain a feature vector containing GC content, predicted transcription factor binding site counts and transposable element coverage. Feature selection is performed to identify the top 100 cell-type-specific features based on a Random Forest (RF) model’s feature importance. Those features selected are then used as input to SACSANN. SACSANN is a stack of two fully connected artificial neural networks, named the Intermediate (IN) and Smoothing (SN) networks, respectively. The IN predicts the probability of an individual bin being in the A compartment, based solely on that bin’s feature vector. The SN takes as input multiple consecutive IN predictions to provide a smoothed prediction of A/B compartment annotation. On the *bottom left,* compartment annotations and predictions for mouse embryonic stem cells chromosome 3 are shown (prediction errors are represented in red).

The entire SACSANN model building procedure (feature selection and training/evaluation) was applied to several A/B compartment annotations derived from neuronal differentiation Hi-C data. For mouse, we used data published by Dixon et al. (2012) (2), Fraser et al. (2015) (11) and Bonev et al. (2018) (12), where the latter two datasets consist of a complete mouse neuronal differentiation. In human, we used hESC Hi-C data from Dixon et al. (2012) (2). When available, both the author’s compartment annotations and those produced by HOMER (13) were used to train and assess models independently. We note that we (and the community) treat compartment annotation as binary (each genomic bin is assigned to either the A or B compartment) and there exist transitional regions whose assignment to A or B is unclear.

SACSANN proved to be accurate across all eight data sets that were tested (average Area Under the Curve or AUC score > 80%, see Fig. 2), which indicates that chromosome compartmentalization is at least partially determined by the underlying DNA sequence. Model performance is similar across chromosomes (Sup. Fig. S1). Notably, SACSANN more accurately predicts A/B compartment annotations produced by HOMER (average AUC score > 88%) than Fraser et al. (2015) (11) compartment calls in mESC. Therefore, we decided to focus on HOMER A/B compartment annotations for further analyses.

**Fig. 2.**
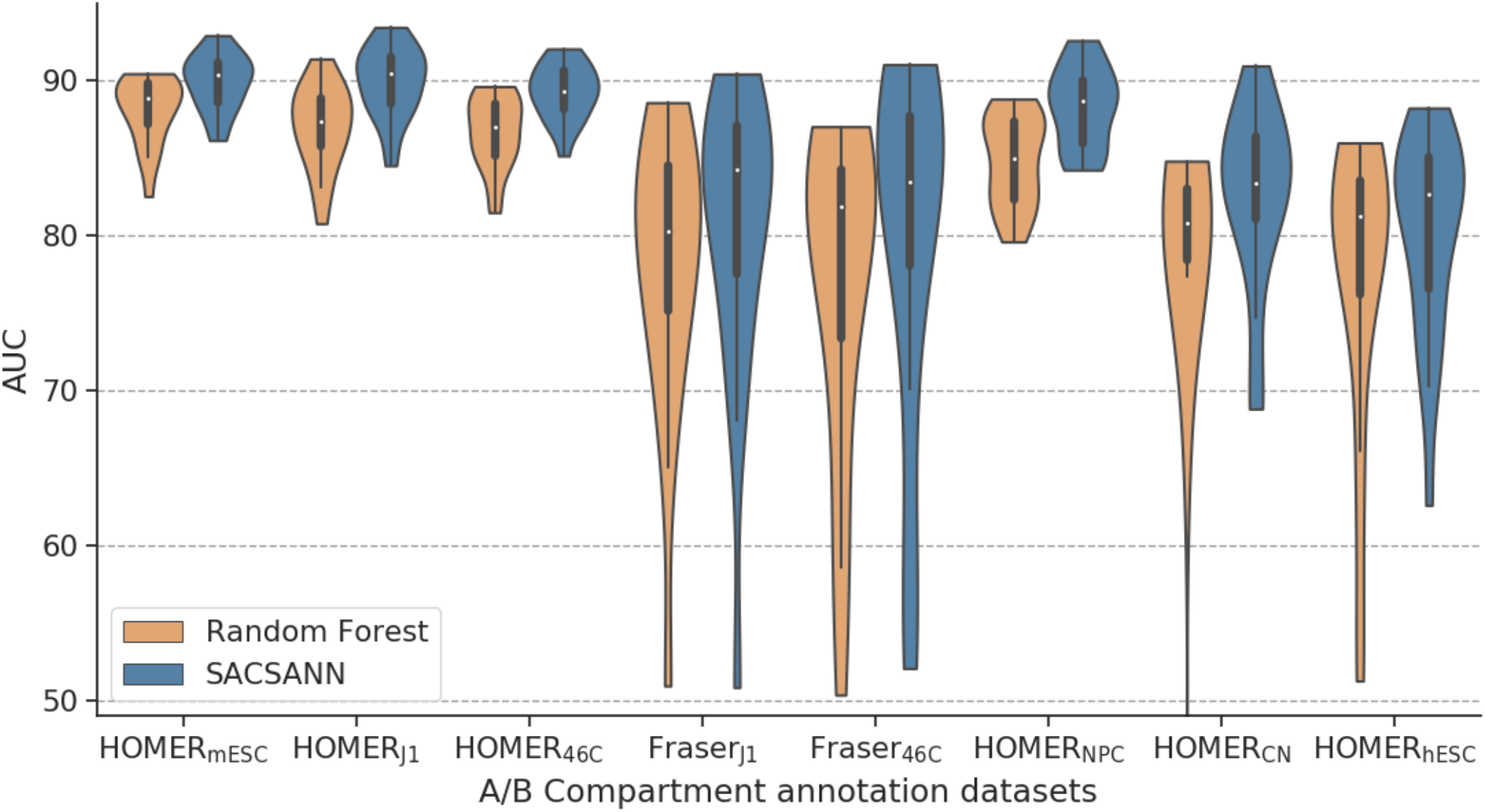
A/B compartments can be predicted from sequence-based features. Accuracy evaluation of SACSANN across multiple different compartment annotations (100 kb resolution), cell types, and species compared against a random forest algorithm. Each violin represents the Area Under the Curve (AUC) score distribution obtained by performing chromosome-wise, leave-one-out cross validation for each datasets (i.e., each violin contains 19 or 22 autosome AUC scores for mouse and human, respectively). Hi-C based compartment annotations were obtained using either HOMER or based on the annotation provided by Fraser et al. (2015) (only for J1 and 46C cells).

Although HOMER’s compartment annotation is binary, it is actually based on the value of the first principal component (PC1), whose magnitude relates to the clarity of the compartment annotation. Bins where the PC1 is close to 0 correspond to regions of poorly defined compartmentalization, and are often found at transitional bins between compartments A and B. Indeed, bins where HOMER and SACSANN disagree tends to have much lower PC1 values than those where they agree (Sup. Fig. S2) suggesting that many of the “errors” found in SACSANN predictions may actually be caused by an uncertain HOMER annotation.

### Compartment annotations are supported by biological evidence

To study the properties of regions correctly and incorrectly predicted by SACSANN, we labeled each bin as either *A* → *A* (HOMER and SACSANN agree to assign the bin to compartment A), *B* → *B* (HOMER and SACSANN agree to assign the bin to compartment B), *A* → *B* (HOMER labels the bin as A compartment but SACSANN predicts it as B), and iv) *B* → *A* (HOMER labels the bin as B compartment but SACSANN predicts it as A). Focusing on mESC (for which epigenetic data is very rich), each genomic bin was associated with an epigenetic/expression state vector describing its overall levels of expression from RNA-seq, histone modifications, and chromatin accessibility (see Methods). These vectors were then hierarchically clustered, revealing three main clusters (Fig. 3). Clusters 1.1 and 1.2, primarily containing bins assigned by HOMER to the A compartment, show high enrichment in H3K36me3, H3K9ac, H3K27ac, H3K4me3, H3Kme3 and CTCF, with high values of DNase hypersensitivity and gene expression. Indeed, A-compartments were previously reported to correlate positively with active histone marks (H3K36me3, H3K4me1 and H3K27ac), open chromatin/euchromatin and highly expressed genomic regions (1, 5). Cluster 1.1 is composed of *A* → *A* bins at 97%, whereas cluster 1.2 is composed of 86% such bins. In comparison, Cluster 2 is enriched for bins assigned by HOMER to the B compartment and depleted for these marks, which is consistent with the trends observed previously. Cluster 2 is composed of 76% *B* → *B* bins.

**Fig. 3.**
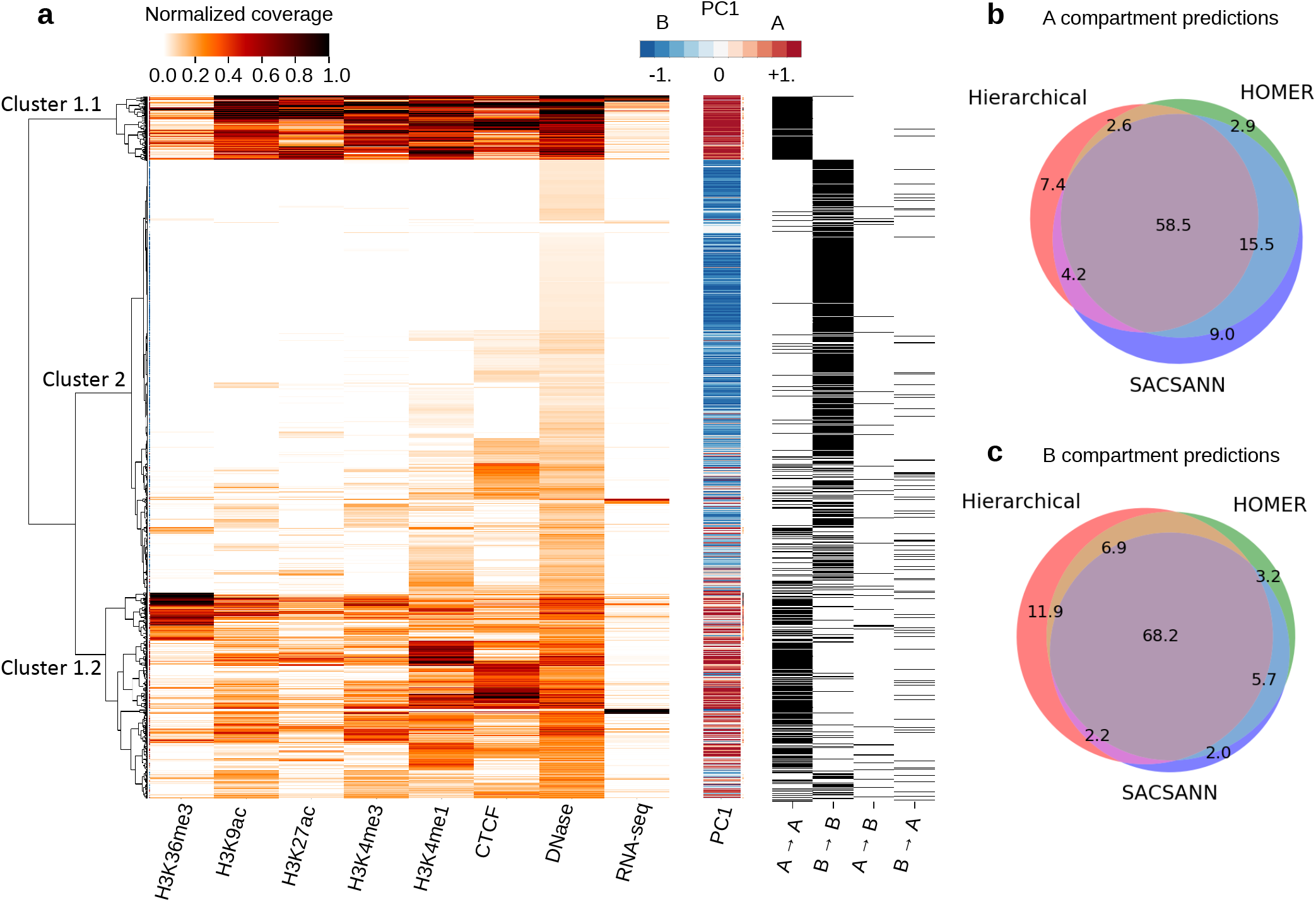
External validation of SACSANN’s compartment annotations. **a**) Hierarchical clustering of the epigenetic/expression state vectors in mESC (23,964 100kb bins in total). Each biological track’s values was scaled between 0.0 and 1.0 with clipping of the top and bottom 1% for visibility purposes. On the *right* of the hierarchical clustering heatmap (in blue to red palette): PC1 value for the corresponding bin as provided by HOMER. *Right* (in black or white), each bin is classified as one of four distinct categories: bins annotated as A by both HOMER and SACSANN (*A → A*), B by both HOMER and SACSANN (*B → B*), A by HOMER and B by SACSANN *(A → B*) or B by HOMER and A by SACSANN (*B* → *A*). **b**) and **c**) Venn diagrams comparing the three compartment annotation methods (epigenetic/expression state vectors clustering, Hi-C based annotations and SACSANN annotations). The diagrams represent the congruence in labelling a bin as being in the A (**b**) or B (**c**) compartment, with respect to the total number of bins labeled for a given compartment type.

Rao et al. (2014) (5) defined a further partitioning of A/B compartments into six sub-compartment types (A1, A2, B1, B2, B3 and B4) for the GM12878 human lymphoblastoid cell line, where each sub-compartment was found to have its own genetic and epigenetic characteristics. To see if this distribution could be replicated with the previous hierarchical clustering, we studied the sub-clusters of Fig. 3a. Clusters 1.1 and 1.2 were found to be analogs of Rao’s sub-compartment A1 and A2 respectively, and differ in their level of enrichment for the histone marks H3K9ac, H3K27ac, H3K4me3, and H3K4me1. However, we were not able to retrieve a similar separation in cluster 2. This might be in part due to the resolution at which this study was performed (100kb against 1kb found in (5)) and cell type differences (mESC vs. GM12878). In addition, the eight available mESC epigenetic data tracks used for clustering is limited compared to the 20 markers used by Rao et al. (2014) (5).

To further compare the A/B compartment annotations made by HOMER and SACSANN, we interpreted the epigenetic state vector clustering (Fig. 3a) as being a third A/B type of compartment annotation method (Fig. 3b), similarly to the reasoning in Fortin et al. (2015) (7). 78% of the genomic bins are found to be annotated consistently by the three methods. The remaining 22% are almost evenly distributed in the other domains of the Venn diagrams, which suggests that no particular method outperforms the others for these genomic loci.

### Compartment establishment rules are transferable across chromosomes

The organizational principles that guide compartment formation still remain unclear and we hypothesize that these principles are broadly shared across chromosomes. So far in this study, SACSANN models have been trained using traditional leave-one-out chromosome-wise cross-validation. To address our hypothesis, we trained SACSANN on individual chromosomes and then predicted A/B compartments for the remaining *n* – 1 chromosomes. Despite a significant decrease in the quantity of training data, the resulting AUC scores remain surprisingly high (89.8% vs 89.9% for mESC, 86.8% vs 88.3% for NPC and 76.0% vs 82.5% for CN (Fig. 4)). Based on these results, we postulate that similar compartment formation rules are shared across chromosomes.

**Fig. 4.**
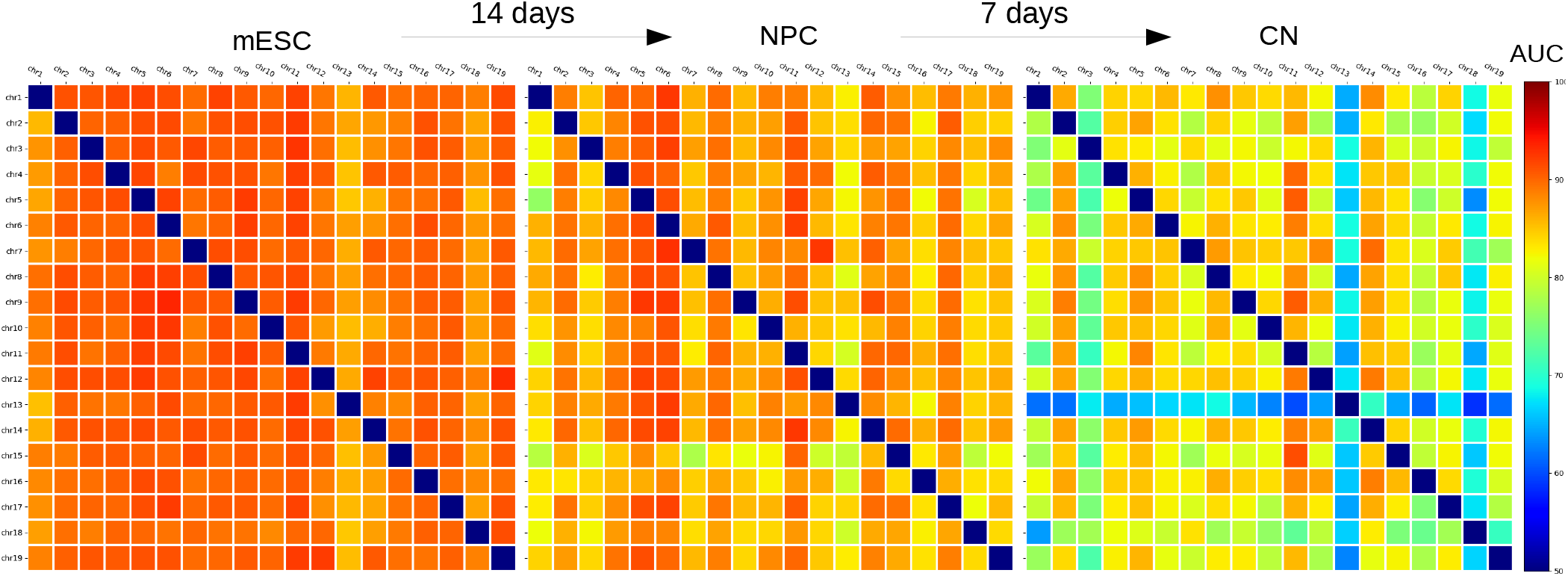
Compartments can be learned from an individual chromosome in mouse neural differentiation. From *left* to *right*: Area Under the Curve (AUC) score heatmaps for SACSANN compartment predictions of mouse Embryonic Stem Cells (mESC), Neuron Progenitors (NPC) and Cortical Neurons (CN) from the HI-C data of Bonev et al. (2018). Each entry in a heatmap is the AUC score achieved by SACSANN when trained and evaluated on the corresponding row and column chromosomes, respectively. The diagonal is set intentionally to an AUC score of 50% as models were not trained and tested on the same chromosome.

In mESC, SACSANN models trained on individual chromosomes provide accurate predictions, although certain chromosomes (e.g., chr1 and chr13) are found to be slightly harder to annotate. As neuronal differentiation proceeds toward NPC and CN, overall prediction accuracy decreases, and certain chromosomes become less useful as training data or harder to predict on. For example, in CN, chr13 becomes both a poor training data set and is poorly predicted from models trained on other chromosomes. We were unable to identify specific characteristics that may explain this phenomenon. Other chromosomes (e.g., chr3 and chr18) become harder to predict but remain relatively useful as training data. One hypothesis for this observation is that SACSANN is able to learn global A/B compartment rules from individual chromosomes, but certain chromosomes (like chr3 and chr18) rely on additional chromosome-specific rules. We relate this hypothesis of chromosome-specific rules to the finding by Rao et al. (2014) of sub-compartment B4, present only on human chr19. We also observe that the corresponding chr19 is poorly predicted in hESC (Sup. Fig. S3). Zhang et al. (2018) (14) have also observed similarly poor predictive capability for chr19 as training data for the prediction of Hi-C contacts in five distinct human cell types.

### SACSANN learns cell-type specific compartmentalization

Genome compartmentalization is cell-type specific to a certain degree (1, 4, 12), although different cell types have the same genome. Different SACSANN models are trained for each cell type, enabling models to vary in the way they interpret genomic sequence to make compartment prediction. To understand the differences between models trained on different cell types, we studied the extent to which a model trained on one cell type is applicable to another. Sup. Fig. S4a shows that a predictor trained on a given cell type indeed achieves the highest AUC score when tested on that same cell type, suggesting that SACSANN is able to learn some cell-type specific compartment properties.

SACSANN’s accuracy was shown to decrease over neuronal differentiation (Fig. 4 and Sup. Fig. S4a). Accuracy is very high for bins whose compartment does not change during differentiation, but lower for those that do (Sup. Fig. S5c). This decrease can be attributed to several factors. First, bins whose compartment changes during differentiation are relatively few compared to those that do not (only 24% of bins change compartment type at least once; Sup. Fig. S5a, providing less training data for SACSANN to learn cell-type specific rules. Second, those bins have less well defined compartment membership (PC1 values close to zero), making HOMER’s annotation less reliable (Sup. Fig. S5c). Third, it may be that the rules that relate sequence to a compartment change are more complex than SACSANN can learn from the limited amount of training data available. Notably, SACSANN’s accuracy is at its lowest for genomic bins that change from the B to the A compartment and is marginally better for bins that evolve from A to B. Nonetheless, it is important to note that the overall AUC score still remains above 80% for all cell types explored.

#### Sequence determinants of compartment predictions

In an effort to understand SACSANN’s use of input features to predict A/B compartments and the differences between celltype–specific predictions, each feature was correlated against SACSANN’s predicted probability of a genomic bin residing within the A compartment. Since GC-content is a major determinant of compartments and a covariate of many Transcription Factor Binding Site (TFBS) counts, we performed the analysis controlling for GC content, using Partial Correlation Scores (PCS) (see Methods). It should be noted that this analysis does not directly interpret how features are used in a SACSANN model but only how individual features correlate to SACSANN’s predictions. PCS of each feature is seen to evolve across differentiation (Fig. 5), providing insight into how sequence determinants correlate with compartment predictions across differentiation. For example, the TFBSs of Nanog, Oct4 and Sox2 are observed to negatively correlate with compartment A prediction scores, with the former two exhibiting progressively reduced importance during differentiation and Sox2 remaining strongly negatively associated throughout. These three transcription factors are critical for the maintenance of pluripotency in mESCs (15–18). The fact that Nanog and Oct4 lose their feature importance over the differentiation is consistent with this maintenance function. In comparison, Sox2 is a determinant in neuron progenitors (19), possibly explaining why it remains of high importance in NPC and CN.

**Fig. 5.**
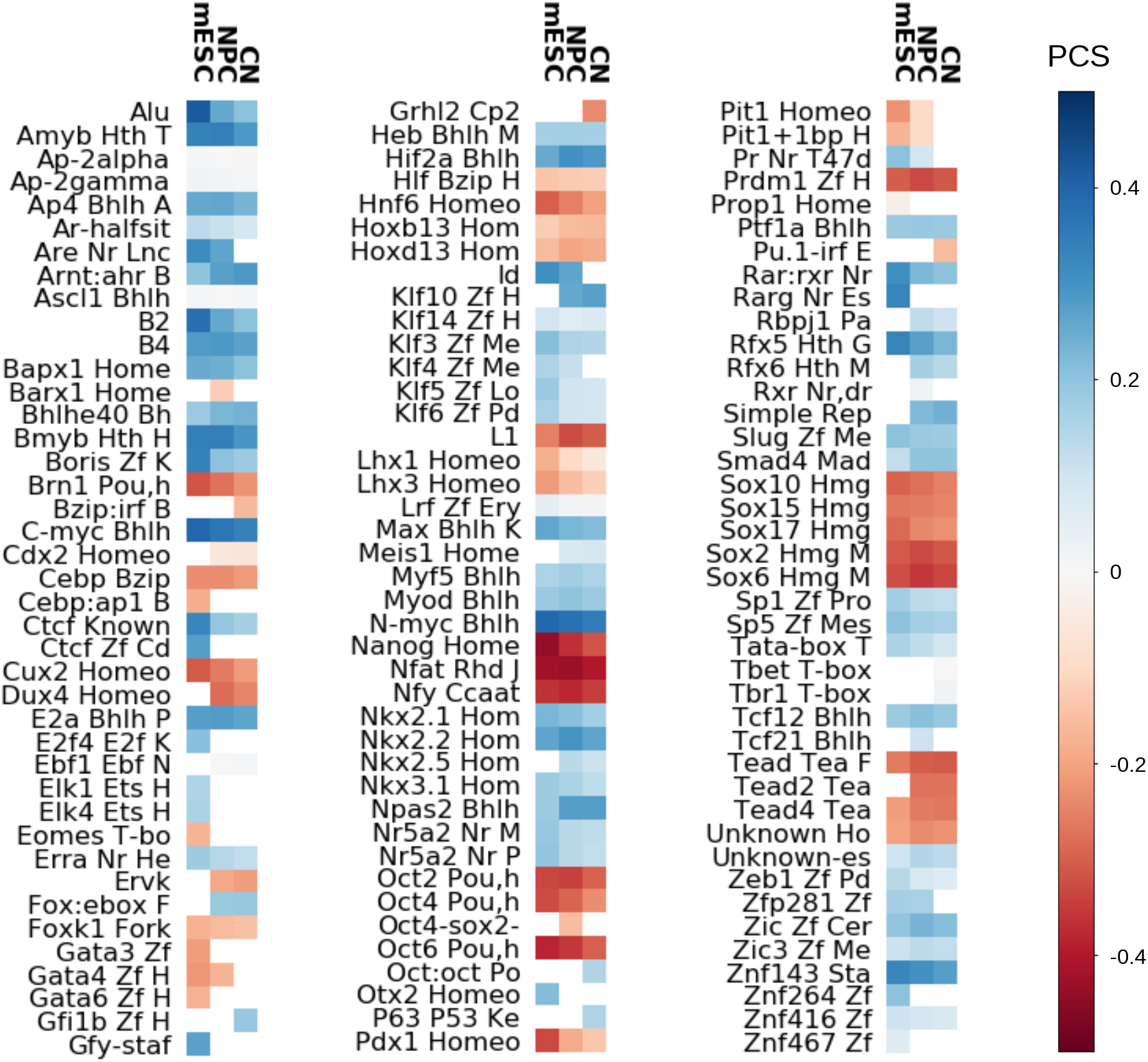
Sequence determinants of chromosomal compartments evolve across mouse neural differentiation. Partial Correlation Scores (PCS) for the top 100 features selected in each cell type for mouse Embryonic Stem Cells (mESC), Neuron Progenitors (NPC) and Cortical Neurons (CN). PCSs correlate feature values with SACSANN’s prediction of a genomic bin being in the A compartment, while controlling for GC content. White entries correspond to features that were not selected to be in the top 100 features used by the corresponding cell-type-specific model. See Sup. File S1 for PCS values and full feature names.

Other features are observed to have their PCS vary across differentiation. This increase is exemplified by the L1 transposable element and TEAD transcription factor, which become more negatively correlated with A compartment annotations over the differentiation. In contrast, TFBSs like HIF2a, bHLHL E40 and Arnt become more highly positively correlated as differentiation occurs. Finally, some features, including Nuclear Factor of Activated T cells and E2A, remain almost constant in their PCS. These observations are consistent with the important role these elements play in brain development (20–22).

### Compartment establishment rules are transferable across species

We next set out to study the extent to which compartment establishment rules could be transferred across species. Training SACSANN to predict mESC compartments yields a predictor that is only slightly less accurate on hESC than mESC (AUC of 80.8% vs. 90.0%, respectively). Results are similar in the reverse direction, with a SACSANN model trained on human achieving high accuracy on both human and mouse (AUC of 80.2% and 85.8% for hESC and mESC, respectively). The counter-intuitive fact that the human-trained model performs better on mouse than on human is attributed to the higher quality Hi-C data in mouse (higher sequencing coverage, more recent protocols), yielding more accurate HOMER-based compartment annotation. Overall, these results suggest that compartment formation rules are at least partially shared between similar mouse and human cell types, which implies a conservation of the compartment establishment mechanism across species in embryonic stem cells.

We then contrasted the hESC- and mESC-trained SACSANN models by comparing their feature PCS values (Fig. 6a). Overall, there is a strong correlation between the way both predictors use features (Spearman correlation coefficient *ρ* = 0.83, p-value = 1.15 × 10^-16^), which supports the rationale that compartment establishment rules are shared across species.

**Fig. 6.**
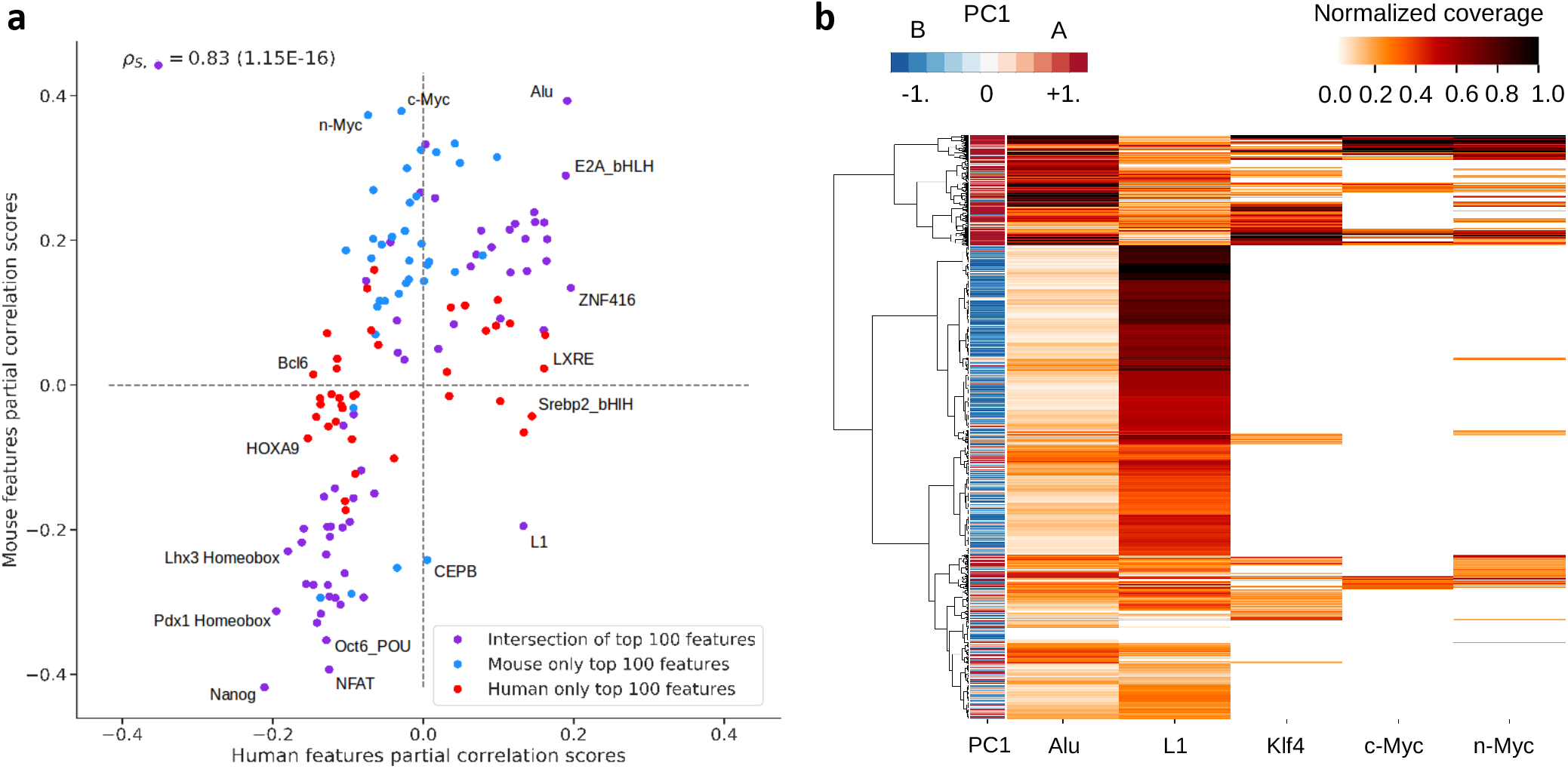
SACSANN identifies sequence determinants of A/B compartments. **a**) For each species (human and mouse), the Partial Correlation Scores (PCS) between each of the top 100 features and A compartment predictions scores by SACSANN in embryonic stem cells are calculated while controlling for GC content. The Spearman correlation score (0.83) of PCSs for the intersection of the top 100 features in human and mouse Embryonic Stem Cells (hESC and mESC, respectively) indicates that a majority of features are found to be similarly used across species by SACSANN. See Sup. File S2 for PCS values. **b**) Hierarchical clustering using a combination of relevant sequence determinants identified by PCS analysis. ChIP-seq peak data tracks were used for transcription factors columns Klf4, c-Myc, and n-Myc. The corresponding PC1 value from HOMER (left) highlights the genome segmentation provided by these new sequence determinants of A/B compartments.

This analysis also reveals that SACSANN predictors rely in part on the presence of various TEs to make their predictions. Notably, the Alu TE has the highest PCS in mESC (0.39) and second highest in hESC (0.19) (note that in mice, Alus refer to the B1 family of TEs, which are SINE elements similar to primate Alus (23)). In humans, Alus are known to be enriched in gene-rich regions (24), which is consistent with the positive correlation to A compartments. Alu elements are known to have a role in regulating the expression of their neighbouring genes in human (25). These elements are also known to have an impact on the primate transcriptome through cis-regulation of RNA editing (26). Surprisingly, LINE-1 (L1) is positively correlated with A compartment predictions in human (0.13), but negatively in mouse (−0.19). In humans, Natale et al. (2018) (24) showed that given their distribution in the genome, L1 and Alu elements represent chromatin regions with opposing features. L1 elements are generally found within AT rich regions of the genome, while Alus prefer gene rich regions. However, since the PCS is calculated by controlling for GC content, this observation is not necessarily contradictory to Natale et al.’s findings. It remains surprising that L1 behaves differently in the two species. SACSANN also relies heavily on a subset of TFBSs, including many homeobox transcription factors, such as Nanog, Oct6, Pdx1 and Lhx3. These factors are found to be negatively correlated with A compartment predictions in both mESC and hESC. Interestingly, Lopes Novo et al. (2016) (27) showed that Nanog is an important regulator of heterochromatin in mESCs.

## Discussion and Conclusion

Chromosome compartments are one of the highest levels of 3D organization and are associated with gene expression (4). Compartment establishment is sequence-encoded and epigenetically-controlled in a cell-type specific manner (7, 8). Yet their sequence determinants remain largely unknown, and to date no attempts had been made to predict compartment organization from sequence alone. In this paper, chromatin A/B compartments were accurately predicted by SACSANN, a ML algorithm using only sequence-based features. SACSANN was shown to be robust across different cell types and species. SACSANN identifies key genomic determinants that define A/B compartments, including Alu TEs and Nanog TFBSs in ESCs. Alu enrichment in the A compartment has been previously linked to a known stabilization role in DNA repair for open chromatin (28). Moreover, SACSANN models trained in one species (mouse or human) proved capable of accurate prediction in the other, suggesting an evolutionary conservation of compartment establishment rules. These conservation results are encouraging for the application of our ML methodology to species where no Hi-C data is currently available, or to computationally inferred ancestral genome sequences (29). The observed similarities in mESC and hESC compartments highlight that such an evolutionary study would probably be insightful given the similar feature behaviors, at least within eutherian mammals.

Improving the cell-type specificity of SACSANN will be addressed in future iterations of the software. The most difficult genomic bins to predict across the mouse neural differentiation (mESC → NPC → CN, see Fig. 4) were those that changed compartment association. To describe these changing bins, a more sophisticated model may be needed along with additional training data and features. One approach to this problem may be to combine data from many different species of a given cell type. This approach is supported by our preliminary results (Fig. 6 and Sup. Fig. S4) that different species share compartment establishment rules. Performance improvements may also be achievable by using predictive models that do not rely on engineered features but instead automatically learn useful data representations, such as convolutional artificial neural networks. Such an approach would not be limited by our current understanding of genomics and could lead to the discovery of new sequencebased determinants, although at a cost in terms of interpretability. Similar ML approaches have been successful at predicting noncoding-variant effects (30) or DNA accessibility (31). Additional performance gains may be achievable through the use of recurrent neural networks (e.g. long shortterm memory networks (32)) to better integrate signals over the entire chromosome length. We envision that the rapid increase in the amount of Hi-C data available to learn from, combined with advances in machine learning will enable further accuracy gains and help more clearly delineate the sequence determinants of compartment formation.

In conclusion, SACSANN allows for the analysis of A/B compartments in species and cell types where Hi-C is unavailable. The establishment of cell-type-specific models for A/B compartment prediction provides valuable insight into the mechanisms at play in establishing chromosomal compartments.

## Methods

### Data sources

The hg19 and mm10 reference genome assemblies were used for human and mouse experiments, respectively. Computational TFBS prediction was performed using HOMER (13) using default parameters, where HOMER’s ‘known_motifs’ collection of vertebrate motifs were applied to reference genome sequences. Repeat Masker (http://repeatmasker.org/) TE annotations were obtained from the UCSC Genome Browser (33).

Sequenced paired-end reads of Hi-C libraries were obtained from published datasets (Table 1) and mapped to their respective reference genomes using the Hi-C User Pipeline (HiCUP) (34).

**Table 1.** Hi-C libraries investigated

| Cell type | Digest enzyme | GEO |
| --- | --- | --- |
| mESC, hESC | HindIII | GSE35156 (2) |
| mESC, NPC, CN | DpnII | GSE96107 (12) |
| mESC, NPC, CN | HindIII, NcoI | GSE59027 (11) |

To produce A/B compartment annotations, both published and HOMER (13) (using default parameters) produced annotation sets at 100 kb resolution were used.

Histone modification, CTCF binding, DNA accessibility, and expression data were obtained Bonev et al. (2017) (12), from the sources listed in Table 2.

**Table 2.** ChIP-seq, DNase-seq, and RNA-seq used from Bonev et al. (2017) (12)

| Name | GEO |
| --- | --- |
| H3K36me3 ChIP-seq | GSM1000109 |
| H3K9ac ChIP-seq | GSM1000127 |
| H3K27ac ChIP-seq | GSM1000099 |
| H3K4me3 ChIP-seq | GSM769008 |
| H3K4me1 ChIP-seq | GSM769009 |
| CTCF TFBS ChIP-seq | GSM918748 |
| DNase Hypersensitivity | GSM1014154 |
| RNA-Seq | GSE96107 |

### Model architecture

Here, we describe the architecture of SACSANN, which consists of two stacked, fully-connected Artificial Neural Networks (ANN) (see Fig. 1 for an overview of the model’s architecture). The first ANN (Intermediate Network [IN]) assigns the probability for a given 100 kb genomic bin to belong to the A compartment. The IN takes as input features representing GC content, TFBS counts and TE counts for the bin (see Feature Selection). All feature values are standardized to have a mean of zero and variance of one.

For training and validation of SACSANN, target values are set to 0 for bins corresponding to compartment A, and 1 for compartment B. The IN is applied separately to each bin of a chromosome. Since compartments have an average size of over 1 Mb and model input/output resolution is 100 kb, most compartments span several consecutive bins. Therefore, a second ANN (Smoothing Network [SN]) is then applied to the output of IN to smooth its A/B compartment predictions. To accomplish annotation smoothing, SN takes as input the output of IN for the current bin *b* and a fixed number of its preceding and succeeding bins: *b* – *w*,…,*b* – 1,*b* + 1,…,*b_w_*. It then produces a revised estimate of the probability of bin *b* belonging to compartment A. The number of neighbors *w* is a tuned hyper-parameter (value selection is discussed below) of the architecture. Overall, IN takes 100 Random Forest (RF) selected, sequence-based features as input. SN observes 1,000 kb windows flanking a given genomic bin *b* to provide the prediction of A compartment assignment, where *w* was set to 10 (i.e., 1 Mb on either side).

Both IN and SN were implemented using the scikit-learn neural_network package (35). Hidden and output layers of the ANNs use sigmoid and softmax activation functions, respectively. Cross-entropy is used as the loss function for both ANNs.

### Model training

SACSANN models were trained on data from an individual or multiple chromosome(s). For multiple chromosome experiments, SACSANN is trained using chromosome-wise leave-one-out cross-validation (i.e., the model is repeatedly trained on all but one of the chromosomes and tested on the left out chromosome). Because bins belonging to A and B compartments are generally not found in equal numbers, we randomly down-sample the majority class in the training set to achieve a balanced representation. The two ANNs composing SACSANN are trained separately, each using the Adam optimizer algorithm (36) and L2 regularization to minimize the cross-entropy loss.

### Hyper-parameter tuning

The Bayesian optimization software Spearmint (37) was used to tune hyper-parameters of SACSANN, which include the number of RF-selected features, initial learning rate, L2 regularization rate, number of hidden layers, number of nodes per hidden layer in each ANN, and number of neighbors to take into account for SN. Parameter tuning was separated into two optimization problems: i) parameter tuning of IN and ii) using hyper-parameters values obtained in i) in combination with Spearmint to tune SN. For the first task, Spearmint was run for 400 iterations with 80 random starts on two mouse Hi-C data sets: mESC from Dixon et al. (2012) (2) and CN from Bonev et al. (2017) (12). For SN, 200 iterations of Spearmint with 40 random starts were performed on the same two Hi-C data sets. For each iteration, 5-fold cross-validation over the entire data set provides an estimate of the current model’s performance. The results of Spearmint are summarized in Sup. File S3. For most parameters, the optimal value found by Spearmint was approximately the same for both Hi-C data sets investigated. For the number of nodes per layer, the nearest power of two to the value outputted by Spearmint was chosen. In the case where parameter values differed across data sets, these parameters were found to not influence the model’s performance greatly and a consensus value was arbitrarily chosen. To validate Spearmint selected values, the resulting SACSANN model was applied to all other Hi-C data sets (see Table 1). Table 3 summarizes the final parameter values of SACSANN’s stacked ANNs.

**Table 3.**
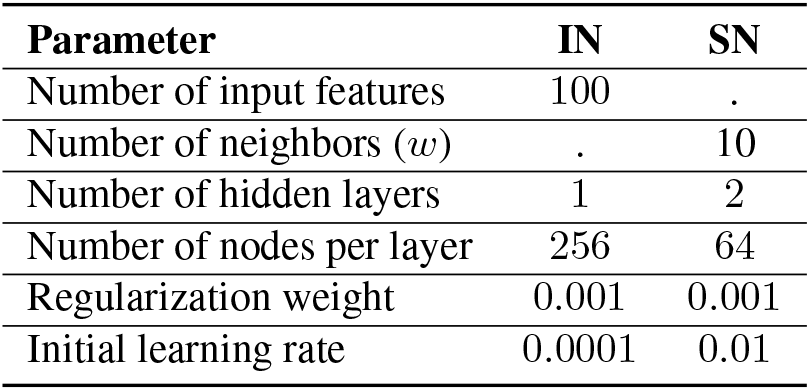
Chosen hyperparameters of SACSANN’s Intermediate (IN) and Smoothing (SN) Networks

| Parameter | IN | SN |
| --- | --- | --- |
| Number of input features | 100 | . |
| Number of neighbors ($w$) | . | 10 |
| Number of hidden layers | 1 | 2 |
| Number of nodes per layer | 256 | 64 |
| Regularization weight | 0.001 | 0.001 |
| Initial learning rate | 0.0001 | 0.01 |

### Feature selection

To avoid the use of redundant features and limit overfitting, mouse and human features (370 and 376, respectively) were ranked according to a RF classifier’s feature importance (implemented in Python’s scikit-learn module (35)). The number of selected features was then tuned as a hyper-parameter of SACSANN using Spearmint.

### Hierarchical clustering

Hierarchical clustering was performed with the Euclidean metric and ‘Ward’ method using the Python module SciPy’s package cluster.hierarchy (38, 39).

### Partial correlation scores

Partial Correlation Scores (PCS) are used to measure the degree of association between each input feature and the probability of a genomic region being in the A compartment. We believe that most computationally-predicted features are driven by GC content and thus, calculate PCSs of these two variables (input feature and A compartment probability) while controlling for GC content. The PCS of two variables *X* and *Y* while controlling for variable *Z* is calculated by correlating the regression residuals of *X* and *Z* against those of *Y* and *Z*. Linear regression is performed between features and GC content. Due to the binary representation of A and B compartment predictions, logistic regression is used to calculate the residuals between compartment predictions and GC content (see Sup. Fig. S8).

## ACKNOWLEDGEMENTS

Authors would like to thank Faizy Ahsan, Emmanuel Bajon, Alexander Butyaev, Samy Coulombe, Josée Dostie, Stefan Kremer, Jacek Majewsky, Vincent Mallet, William Pastor and Zichao Yan for their valuable and constructive feedback during the development of this project and manuscript.

## Supporting information

**S1 File.** Partial Correlation Scores (PCS) and definitions of the abbreviated feature found in Fig. 5.

**S2 File.** PCS values of the abbreviated features found in Fig. 6a.

**S3 File.** Summary of Area Under the ROC Curve (AUC) scores achieved for tested hyperparameter combinations.

Supplemental files are found in the SACSANN GitHub repository: https://github.com/BlanchetteLab/SACSANN/tree/master/supplemental_files

---

**Supplementary Figure S1.**
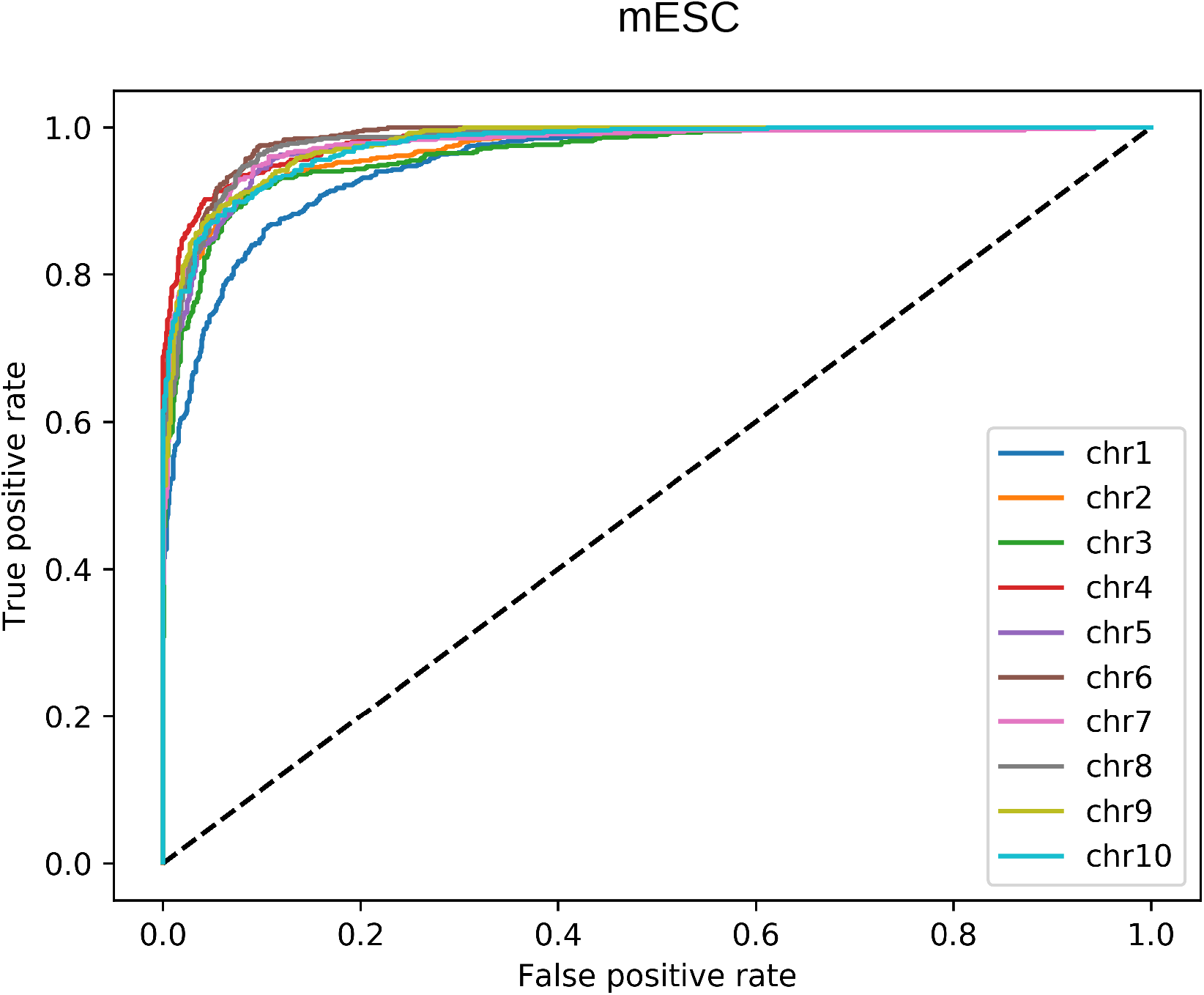
SACSANN accurately predicts chromosome compartments. Example ROC curves of SACSANN predictions for the first 10 chromosomes in mESC.

**Supplementary Figure S2.**
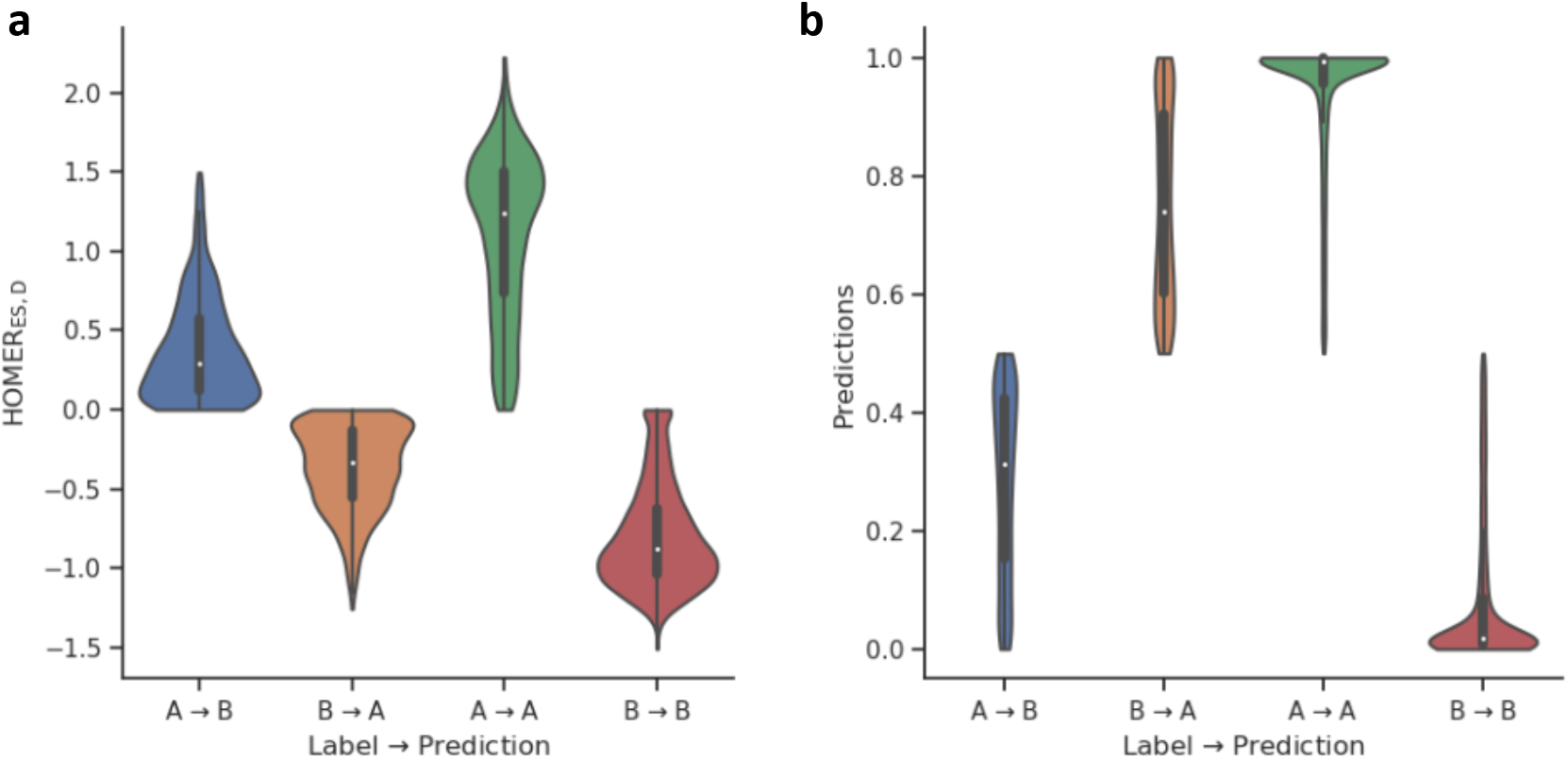
Error Analysis of SACSANN predictions in mESCs. **a**) HOMER PC1 value distribution. Each genomic bin assigned one of the following four distinct classes: 1) compartments annotated as A by both HOMER and SACSANN (*A* → *A*), 2) B by both HOMER and SACSANN (*B* → *B*), 3) A by HOMER and B by SACSANN (*A* → *B*), and 4) B by HOMER and A by SACSANN (*B* → *A*). **b**) SACSANN predicted probability of a genomic bin being found in the A compartment for each compartment class.

**Supplementary Figure S3.**
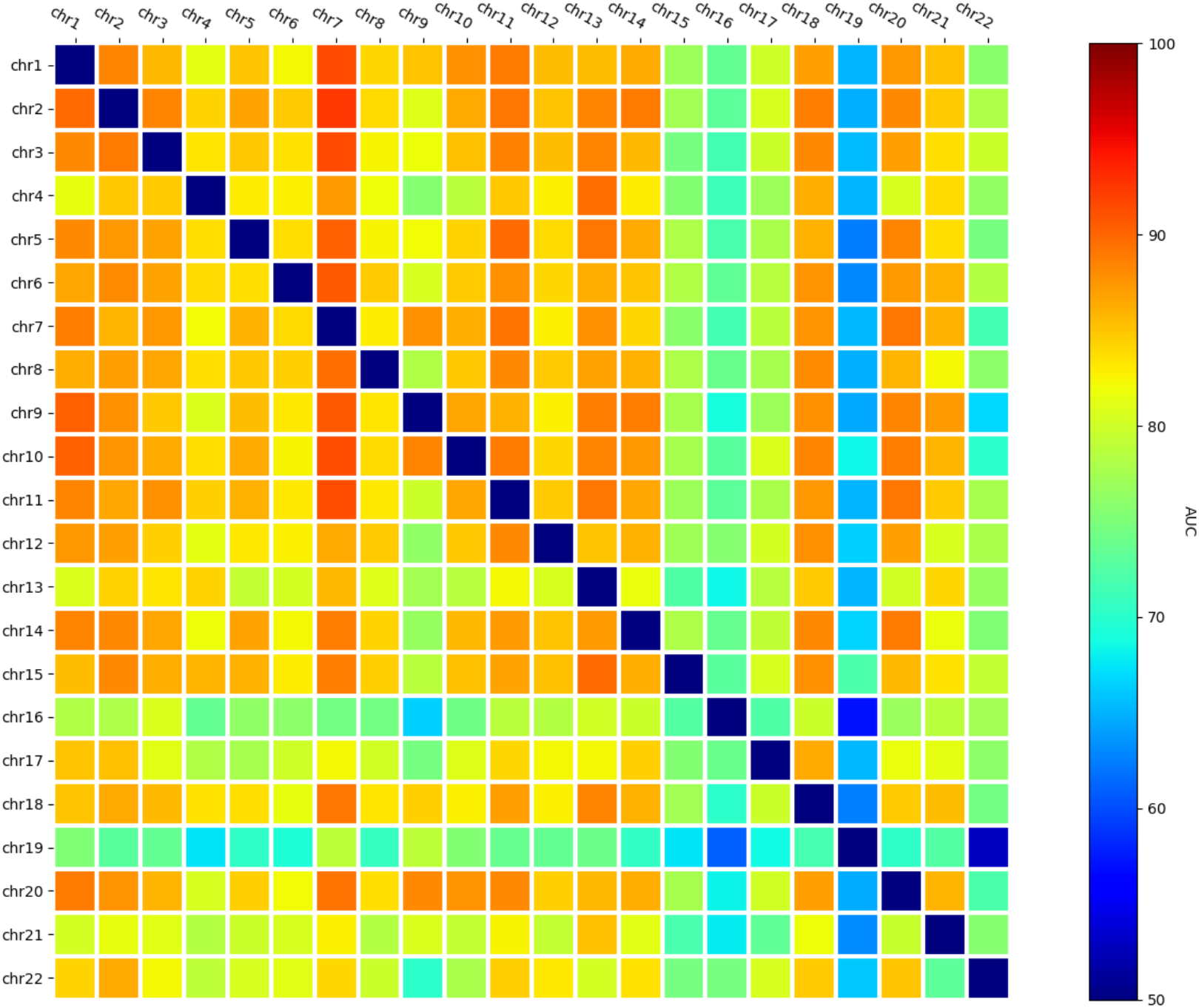
Individual chromosome training for hESCs. Repetition of the analysis found in Fig. 4. See caption of Fig. 4 for a description of the analysis.

**Supplementary Figure S4.**
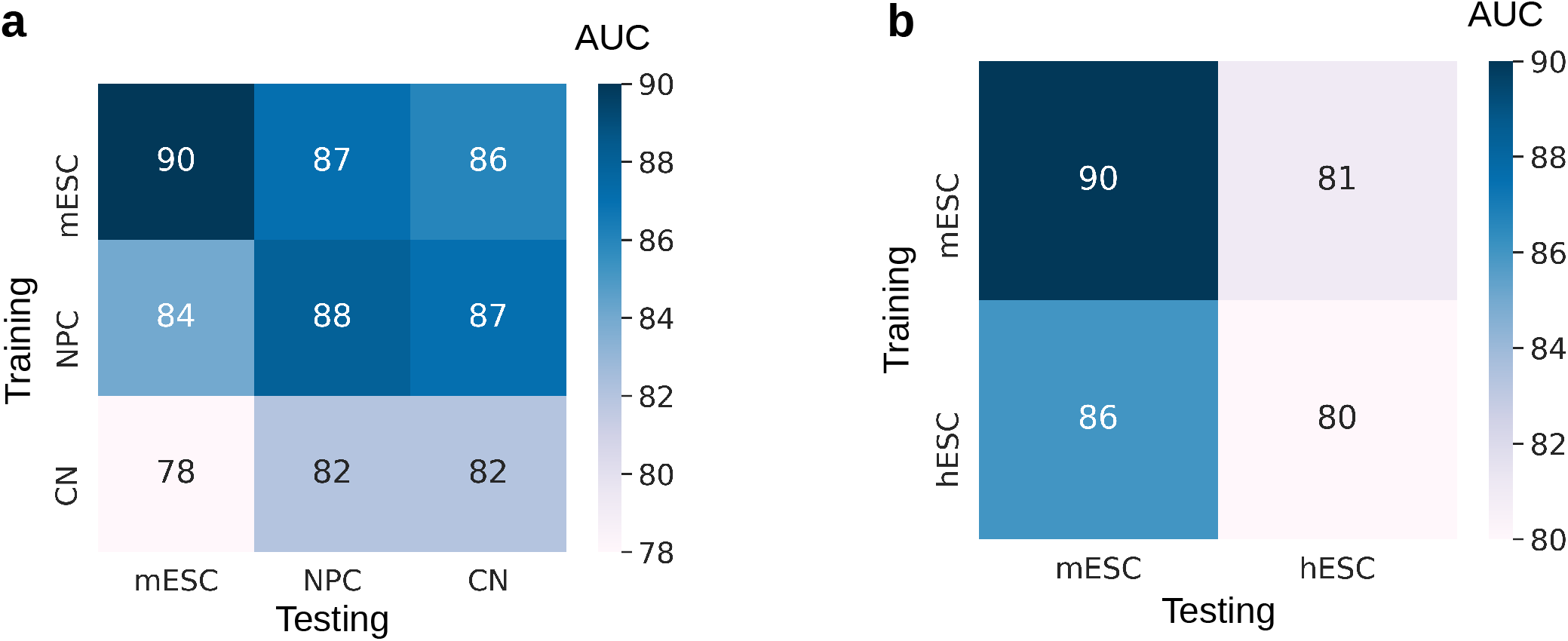
Compartment predictions across cell types and species. Chromosome-wise cross-validation AUC scores for SACSANN predictions of **a**) neural differentiation and **b**) hESC vs. mESC. SACSANN achieves the highest AUC score for the cell type the algorithm was trained on in the neuron differentiation. Interestingly, for the comparison of hESC vs. mESC, SACSANN models trained in mouse are slightly more accurate than those trained in human.

**Supplementary Figure S5.**
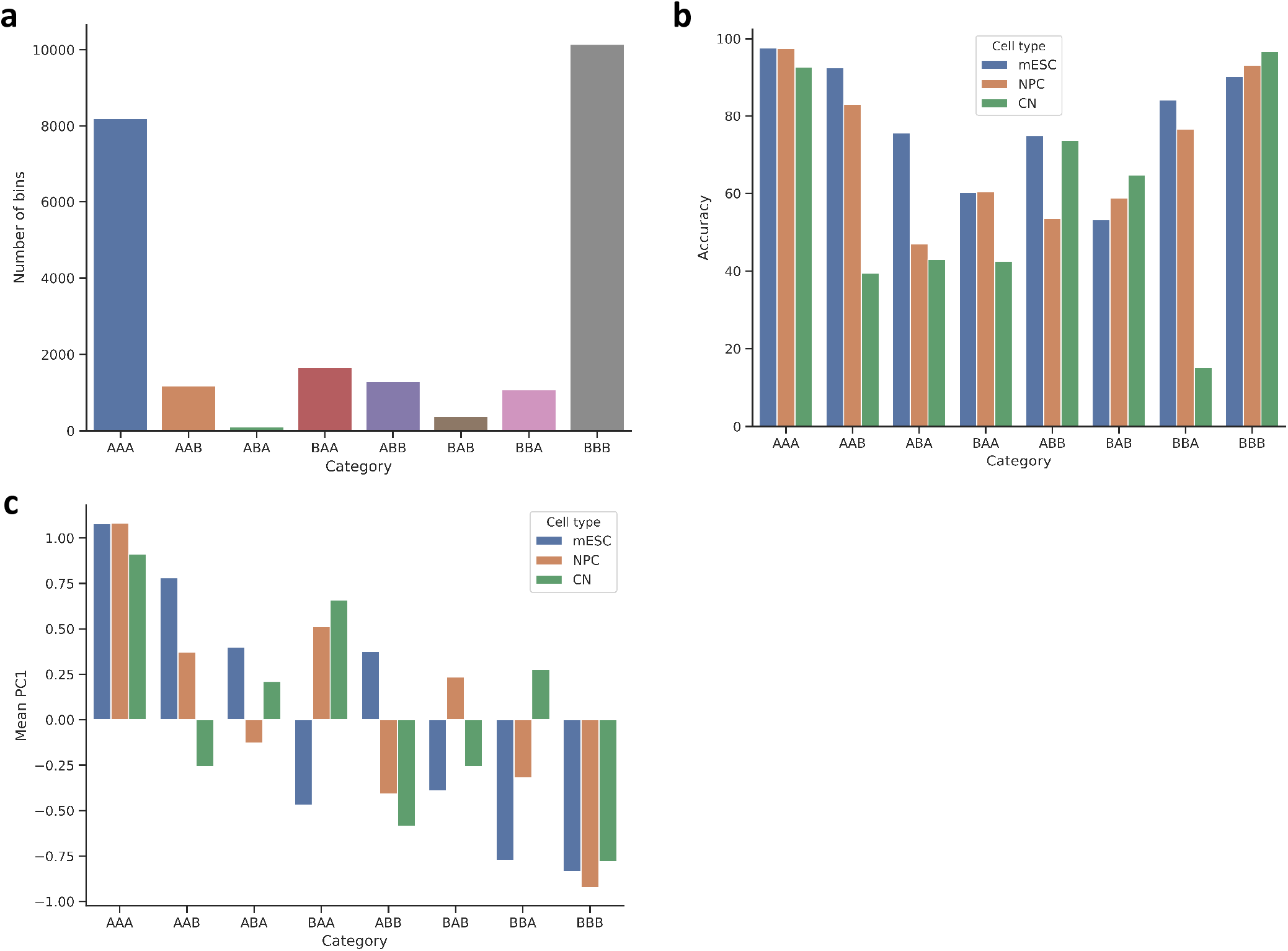
A/B Compartment analysis across mouse neuronal differentiation. **a**) The number of genomic bins as a function of compartment class, where class label ‘XYZ’ is the compartment annotation type for mESC (X), NPC (Y) and CN (Z), respectively. **b**) and **c**) SACSANN accuracy and HOMER PC1 values as a function of compartment and cell type.

**Supplementary Figure S6.**
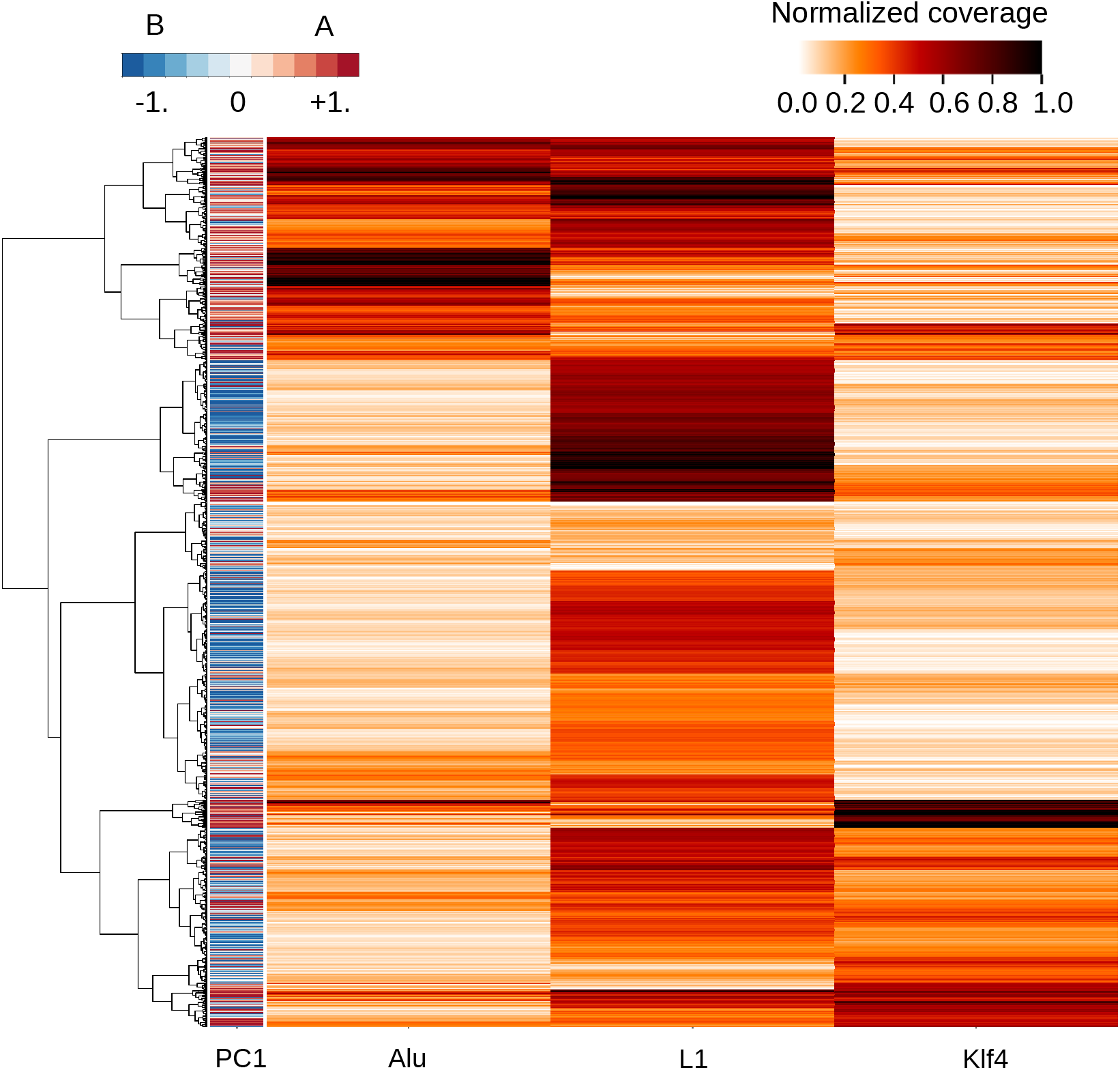
SACSANN identifies sequence determinants in hESCs. Hierarchical clustering of relevant features in hESC according to their PCS score. Klf4 binding sites were obtained from ChIP-seq peak data (see Methods).

**Supplementary Figure S7.**
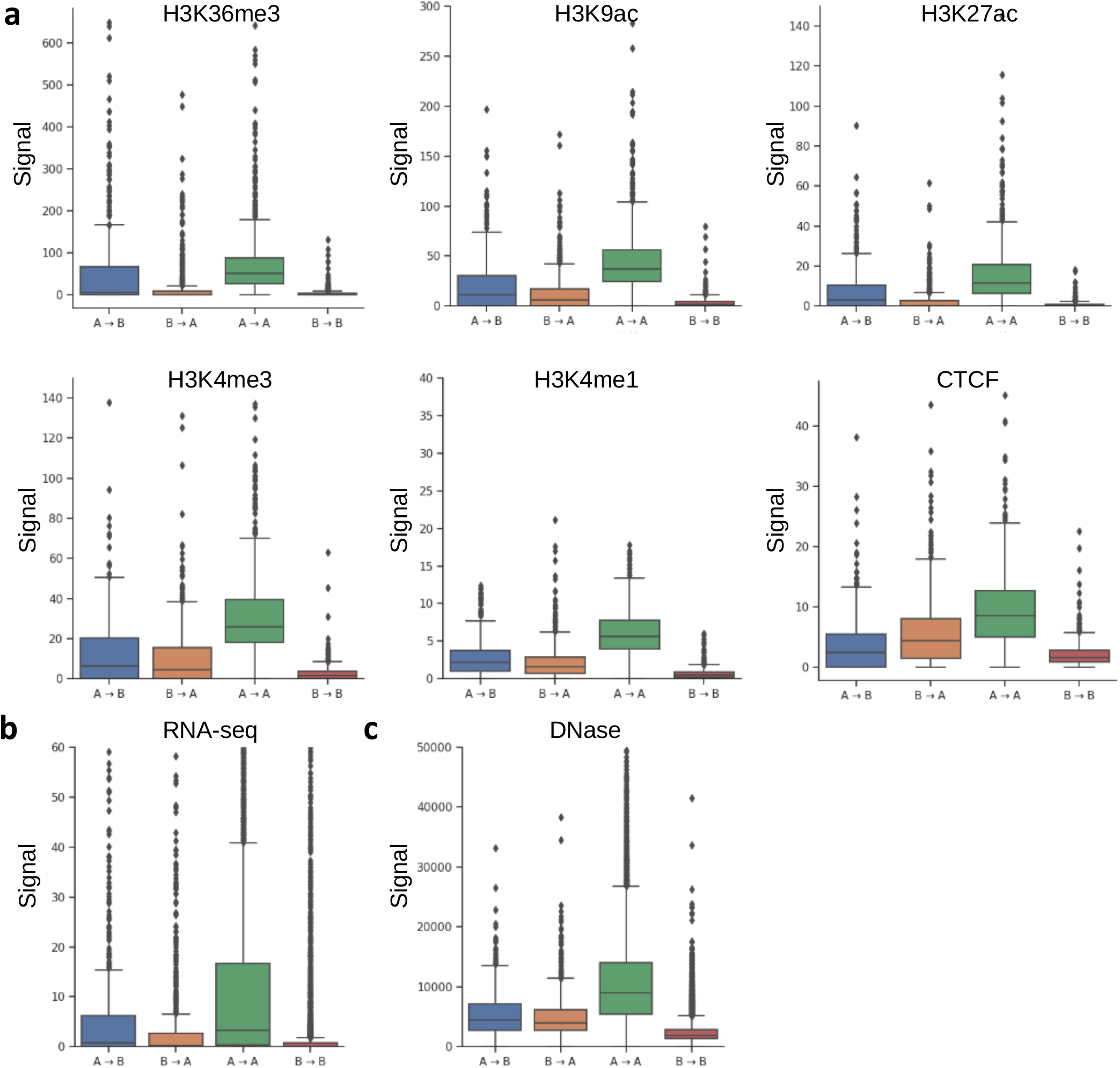
External data re-partition in mESCs. Each data track was binned according to the same categories described in Sup. Fig. S2, where: **a**) ChIP-seq peak data for different histone marks and the CTCF transcription factor, **b**) RNA-seq and **c**) DNase-seq.

**Supplementary Figure S8.**
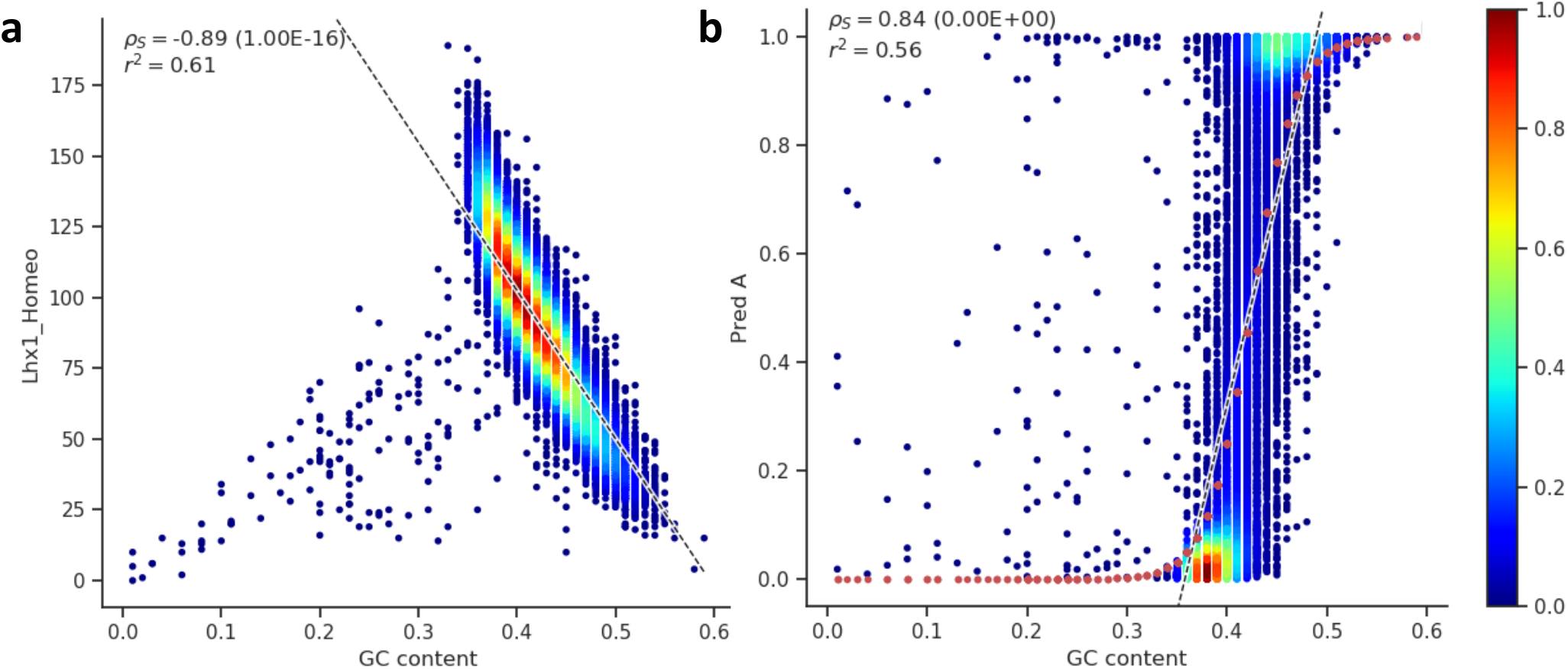
Rationale for linear and logistic regression in PCS. Most features are assumed to be driven by GC content. To address this assumption, we performed Partial Correlation Score (PCS) analyses while controlling for GC content instead of standard correlation analyses. In calculating a PCS, the residuals of two regressions are correlated to quantify the amount of signal not explained by the controlled variable (i.e., GC content). Based on correlations, we chose to apply linear and logistic regressions (black and red dotted curves, respectively) for **a**) features and **b**) SACSANN predictions.

## Notes

### Competing Interest Statement

The authors have declared no competing interest.

